# Discovery of small-molecule inhibitors targeting the ribosomal peptidyl transferase center (PTC) of *M. tuberculosis*

**DOI:** 10.1101/604777

**Authors:** Benjamin Tam, Dror Sherf, Shira Cohen, Sarah Adi Eisdorfer, Moshe Perez, Adam Soffer, Dan Vilenchik, Sabine Ruth Akabayov, Gerhard Wagner, Barak Akabayov

## Abstract

*M. tuberculosis* (*Mtb*) is a pathogenic bacterium that causes tuberculosis, which kills more than 1.5 million people worldwide every year. Strains resistant to available antibiotics pose a significant healthcare problem. The enormous complexity of the ribosome poses a barrier for drug discovery. We have overcome this in a tractable way by using an RNA segment that represents the peptidyl transferase center as a target. By using a novel combination of NMR transverse relaxation times (T_2_) and computational chemistry approaches, we have obtained improved inhibitors of the *Mtb* ribosomal PTC. Two phenylthiazole derivatives were predicted by machine learning models as effective inhibitors, and this was confirmed by their IC_50_ values, which were significantly improved over standard antibiotic drugs.

---

A recent U.K. government review (https://amr-review.org/) on the global threat of anti-microbial resistance predicts that by 2050 2.5 million people will be at risk each year of dying from drug-resistant strains of *Mtb*, the causative agent of tuberculosis (TB). Two crucial stages in the ongoing effort of drug discovery – both of which are addressed in this study – are the selection of the drug target for the specific pathogen and the discovery of lead compounds. One of the major obstacles in pursuing this goal is the complexity of the bacterial ribosome as a target. We have overcome this barrier by targeting an RNA segment of the ribosomal PTC of *Mtb*.

The PTC is a universally conserved ribonucleotide chain that catalyzes the formation of peptide bonds during peptide chain synthesis and constitutes a highly selective drug target site within the ribosome.^5^ Indeed, being the heart of the gene translation machinery, most of the available antibacterial compounds that target the large (50S) ribosome subunit of bacteria act on the PTC.^6, 7^

We have chosen a fragment-based screening (FBS)-virtual screening tandem (Fig. 1A) by virtue of the two major advantages of FBS over high throughput screening: FBS requires a method such as NMR, to detect binding,^8^ and fragment-based libraries cover a larger chemical space.^9^

**Fig. 1.**
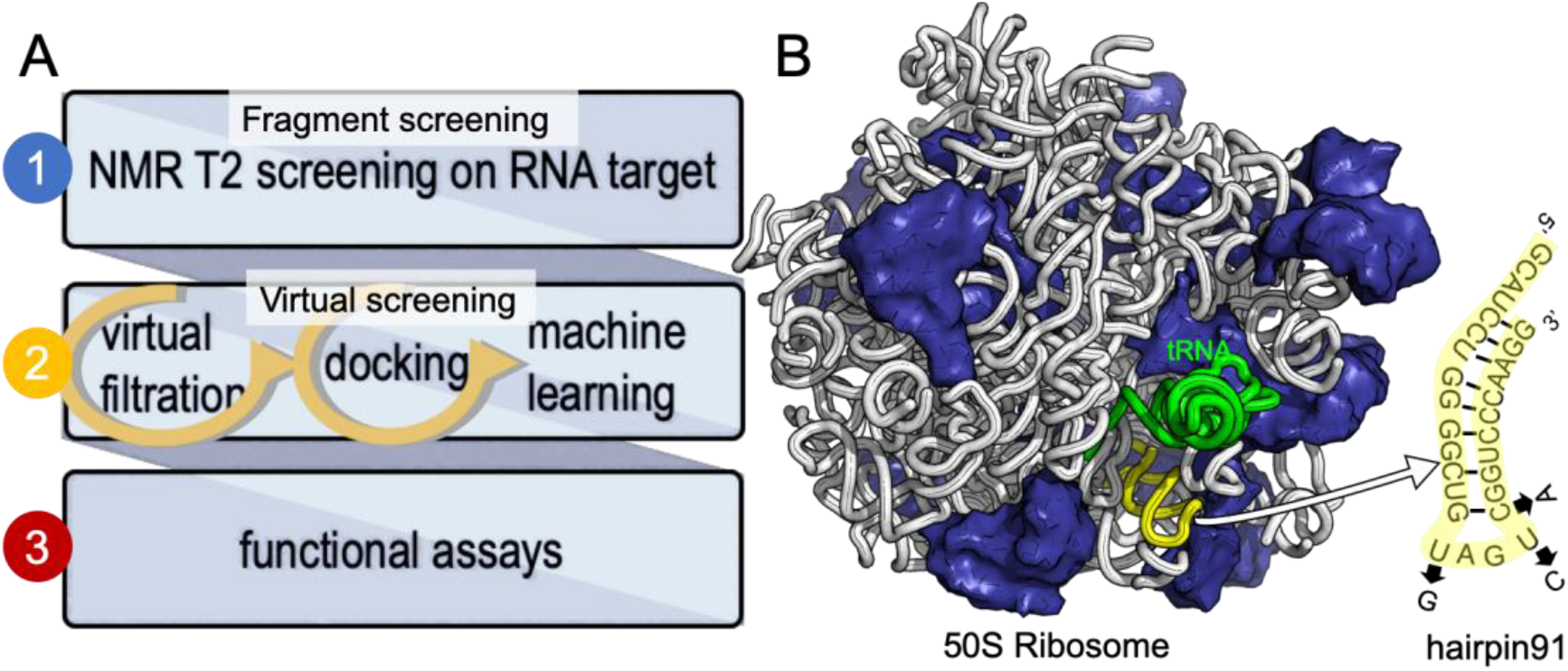
Development of small molecule inhibitors for ribosomal PTC. A) the workflow combines NMR-fragment based screening with virtual screening. Using NMR (T_2_ relaxation) and a fragment library, we identified scaffolds that bind hairpin 91 in the ribosomal PTC of *Mtb.* These scaffolds were used to filter larger compounds containing the fragment molecules from the ZINC database.^1^ Nearly 1000 compounds containing phenylthiazole were docked, using Autodock software,^1, 2^ into the PTC of the available crystal structure of a bacterial ribosome, and hits were ranked on the basis of the binding energy. Conclusions as to structure activity relationships were drawn using machine learning algorithms, and ten compounds were selected and tested for their ability to inhibit translation in *Mtb*. B) The 50S ribosome subunit of *S. aureus* (PDBID: 4WCE^3^) contains two RNA molecules (white) and 34 proteins (blue). An A-site tRNA (PDBiD: 4v4W^4^, green) superposed to the 50S ribosome indicates the PTC. The sequence of RNA hairpin (hairpin 91, yellow) was used for T_2_ relaxation screening. The modified nucleotides in the sequence for *Mtb* are marked in arrows.

We have previously used saturation transfer difference (STD) spectra to identify fragments that bind to DNA primase.^15^ The current study addresses the problem inherent in NMR-saturated difference spectroscopy that the low distribution of protons in nucleic acids (compared to proteins) makes it difficult to excite only the macromolecule and hence to obtain an effective transfer of magnetization to the bound fragment molecule. Here, we used T_2_ relaxation NMR spectroscopy for the initial screening of fragment molecules (Fig. 1A). Using the Carr-Purcell-Meiboom-Gill (CPMG) sequence, we have utilized a property that small molecules bound to larger RNA molecules adopt the relaxation time of the complex.^10, 11^ Obtaining the transverse relaxation rates in the absence and presence of RNA allows determining which fragments bind to RNA. As a target for screening, we used the 29 mer RNA sequence that corresponds to hairpin 91 in the PTC of *Mtb* (Fig. 1B).

After an initial feasibility screen, we performed a full screen with 1000 compounds from the Maybridge Ro3 1000 collection using pools of 9 to 11 compounds per sample. The fragment exhibiting the largest change in T_2_ upon RNA binding is [2-(3-chlorophenyl)-1,3-thiazol-4-yl]methanamine (Fig. 2, Tables S1-2 present the summary of NMR screening results and values for the transverse relaxation time of the best hit, respectively). Out of the nine fragments with the largest changes in T_2_, four contain a phenylthiazole moiety. This moiety was therefore used as the scaffold for the subsequent optimization steps in the FBVS workflow (Fig. 1A).

**Fig. 2.**
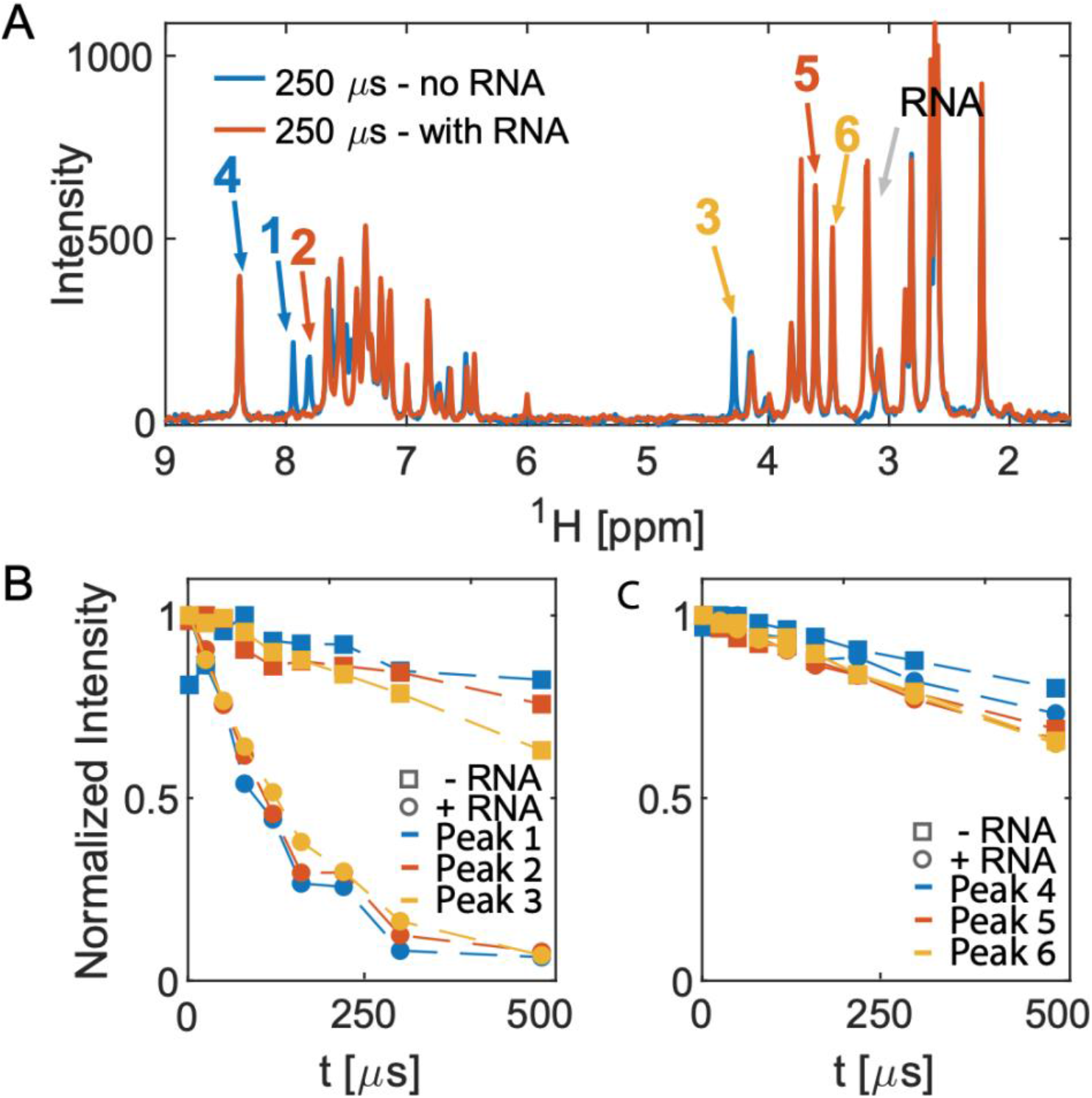
1D CPMG spectra and T_2_ analysis of pool 81. A) Overlay of the two spectra in the absence (blue) and presence (orange) of RNA at the longest T_2_-delay. B) and c) show the intensity of the peaks numbered in A). Peaks 1-3 all originate from one molecule, [2-(3-chlorophenyl)-1,3-thiazol-4-yl]methanamine, which shows the largest change in T_2_ upon addition of RNA. The other three peaks, peaks 4-6, originate from different molecules in the same pool do not show any significant change in T_2_ upon addition of RNA.

The next step in the FBVS workflow was to select a subset of phenyl thiazol-containing molecules from a library of 230M compounds in the available ZINC database.^1^ This step yielded 919 molecules which were docked against the available crystal structure of the ribosomal PTC and were ranked based on their binding energy (large −ΔG, Table S3). We exploited the available structure for the large ribosomal subunit of *S. aureus*^3^ (Fig. 1B) to extract the PTC for docking; the high sequence conservation between the PTCs of *Mtb* and *S. aureus* (>85% for the PTC and 90% for hairpin 91), allows the use of the ribosomal PTC of *S. aureus* as a proxy for the target in *Mtb*.

The inhibition effect of ten compounds with the highest BOND values (**1-10**, Fig. 3A) on *M. smegmatis* ribosomes was tested in a bacterial coupled transcription-translation assay (*see* Methods^12^) in which the expression of luciferase gene was measured. *M. smegmatis* ribosomes are often used to test antitubercular agent, as for the ribosomal PTC used in this study (U2673-C2836), there is 100% identity between *Mtb* and *M. smegmatis* (the identity of the whole 23S rRNA is 2873/3150, 91%). A reconstituted mycobacterial translation apparatus was obtained as described previously.^13^ The results indicated effective inhibition of *M. smegmatis* ribosomes by molecules **2** and **8**, both containing piperazine in their R^1^ position (Fig. 3B). Notably, the IC_50_ values for molecules **2** and **8** were 9.1 and 2.8 μM, respectively. These values are superior to the IC_50_ for chloramphenicol (7.1 μM), a broad-spectrum antibiotic that binds the PTC, thereby inhibiting ribosome activity (Fig. 3B, Table S4).

**Fig. 3.**
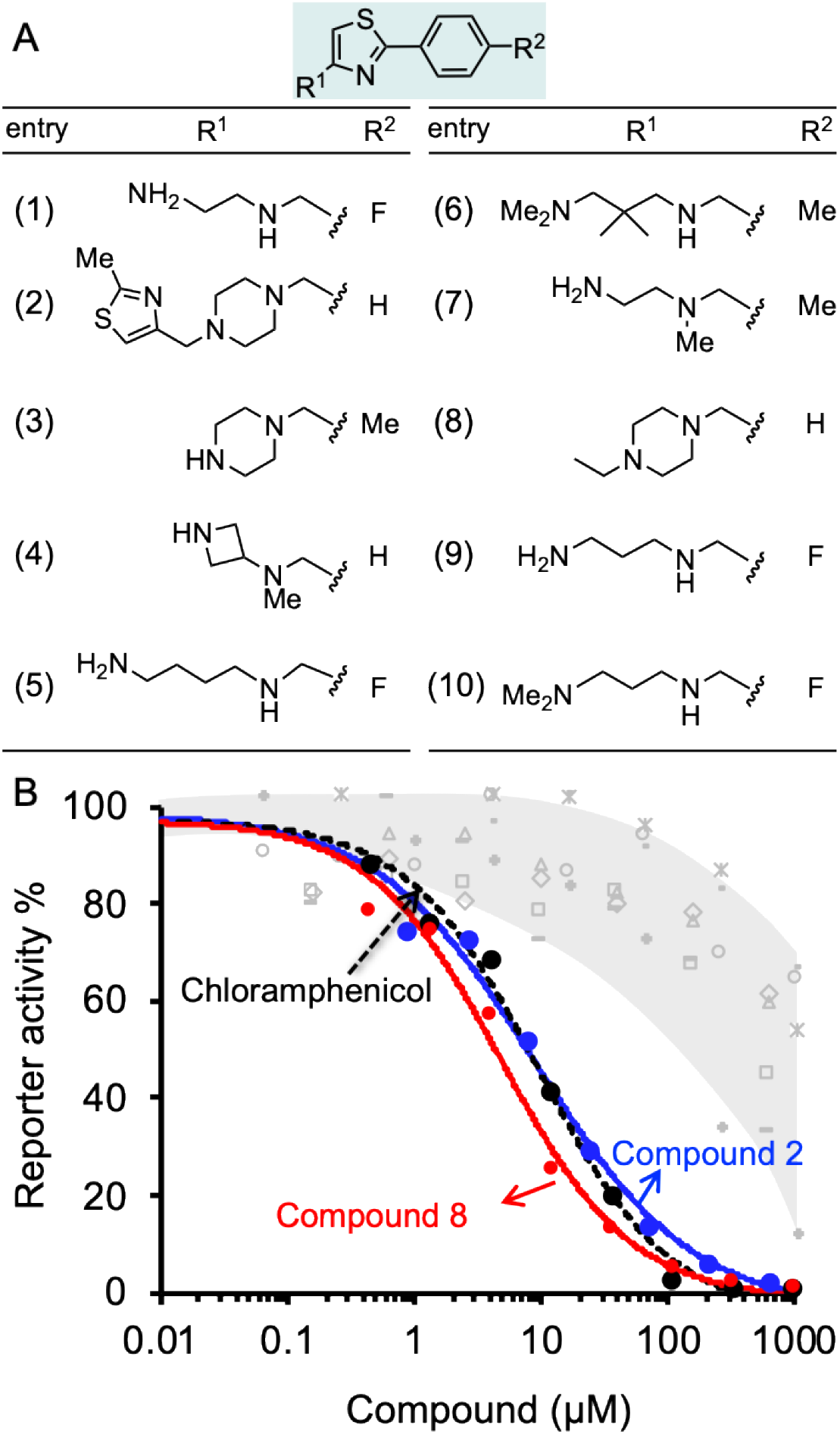
Effect of compounds on protein synthesis measured by in vitro translation reaction. A) Structures of ten phenylthiazol-containing molecules with the best docking binding free energy (*Δ*G_*bind*_). B) The luciferase gene was inserted into a plasmid downstream from the T7 RNA polymerase promotor. The translation reaction is described in details in Methods. The reaction buffer and compounds 1-10 were added in concentrations ranging from 0.15 μM to 1 mM and incubated at 37 °C for 1 h. The effect of the compounds was tested against M. *smegmatis* ribosomes. The effect of compounds 2, 8, and chloramphenicol is colored blue, red and black, respectively. The effect of molecules 1, 3-7, and 9-10 is indicated in grey.

To show specific binding of molecules **2** and **8** in hairpin 91 of the ribosomal PTC an in-line probing assay was used. In-line probing assay has been used to elucidate the binding properties of a variety of metabolites and small molecules to RNA molecules.^14^ This assay utilizes a slow “in-line” nucleophilic attack on a phosphodiester bond in RNA by the 2’ oxygen atom of a spatially adjacent nucleoside. This selective cleavage reveals secondary structural features of the backbone RNA when the products of the cleavage are examined using gel electrophoresis. Binding of a small molecule to the RNA “locks" the RNA in an in-line rigid conformation and promotes cleavage of the linkage involved. The outcome is elevated intensities of the gel bands, indicating hotspot linkage in the RNA target. The rigidity of some linkages along the RNA makes others less likely to adopt an in-line conformation, which is indicated by reduced gel bands attributed to the cleavage sites. Such a change in the pattern of the RNA cleavage is observed in an in-line probing gel upon titration of the small molecule inhibitors, **2** and **8** (Fig. 4A-B), thus confirming the binding site of the molecules.

**Fig. 4.**
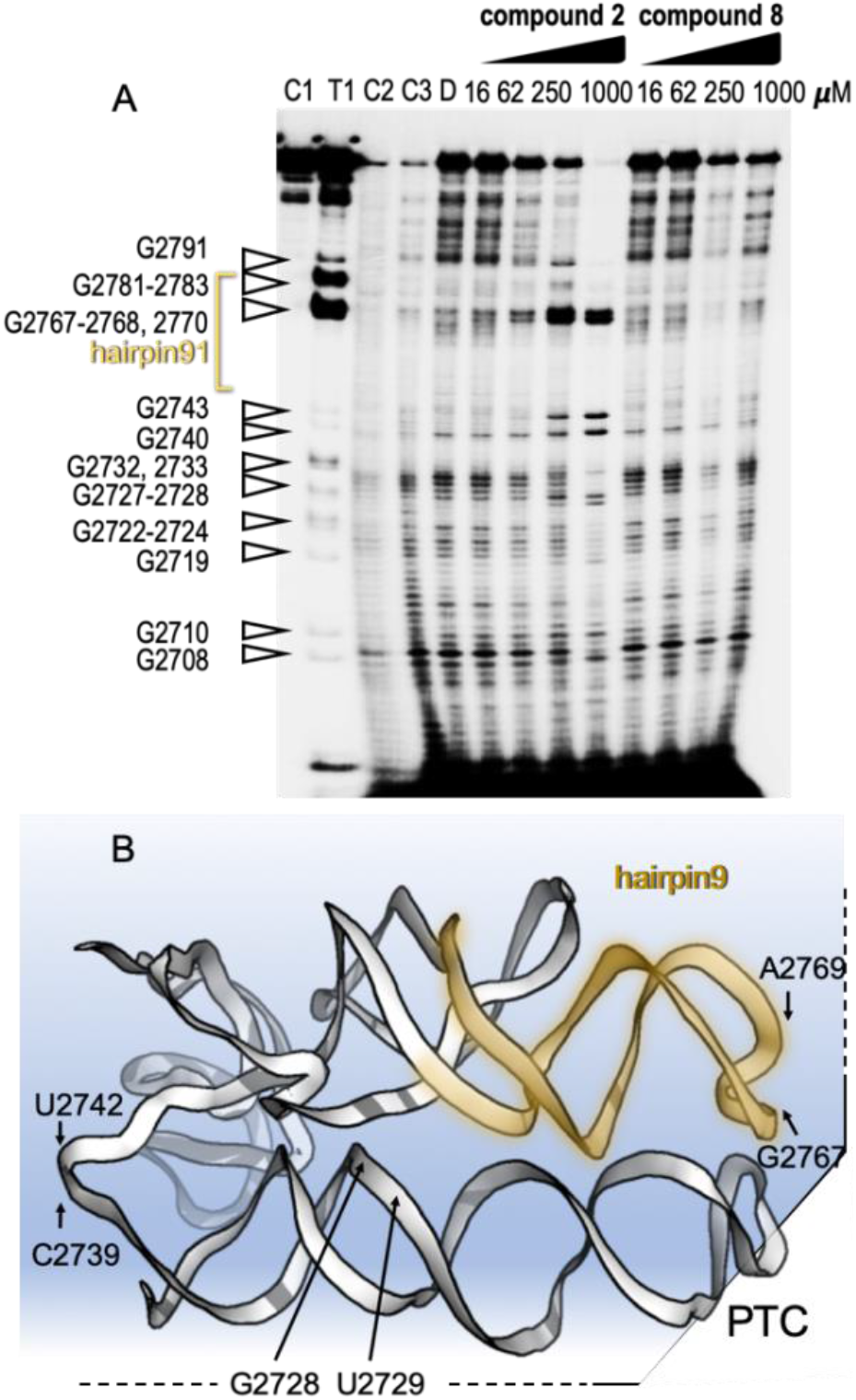
Mechanism of binding of small molecules to PTC. A) In-line probing gel image of ribosomal PTC of *Mtb.* The RNA segment used for this experiment (U2687-C2850) was transcribed *in-vitro* and purified (Fig. S1). The reaction contained 5 nM RNA labeled with ^32^P at its 5′ terminus and in-line reaction buffer. After incubation at 25 °C for 50 h, the RNA product was analyzed by denaturing gel analysis and autoradiography. Right: Lanes 1-3 contain the untreated RNA (C1), partially digested using T1 RNase (T1), alkaline digested (−OH, C2). Lanes 4-5 show the cleavage pattern of the RNA in the absence of the small molecules (C3) and in the presence of DMSO (D). Lanes 6-9 and lanes 10-13 contain compound 2 and compound 8, respectively, in the indicated concentrations. B) Structural model of the RNA shows hotspots of scission of PTC in the presence of compounds 2 and 8.

The docking data was subsequently used for establishing machine learning design principles for drug-sized molecules with increased binding efficiency (BOND) to the ribosomal PTC. Currently, there are no guidelines as to which modifications will be more advantageous. Instead, this step is done by an exhustive brute-force search (medicinal chemistry). Such an approach is not scalable in the face of big-data availability. Machine learning models have been used for a variety of applications in drug discovery.^15–18^ To establish the design principles for drug sized molecules, we trained machine learning models on the docking data. Specifically, a feature engineering step yielded three properties for each of the 11 atom positions in the scaffold (33 features overall). The three properties MOD*i*, DIST*i*, and VAR*i* represent various geometrical features of the phenylthiazole scaffold. We trained two statistical regression models for BOND as a function of MOD*i*, DIST*i*, and VAR*i* – linear regression and random forest^19, 20^. The most critical features, out of the total of 33 features, for predicting BOND were extracted from each machine learning model. Specificially, locations 6 and 8 on the scaffold were found to be significant, and relevant guidelines for the structure to be attached were formed (see Fig. S4). We ran the machine learning models on the 10 molecules, and additional antibiotics. The predicted BOND value for molecule **2** (1-[(2-methylthiazol-4-yl)methyl]-4-[(2-phenylthiazol-4-yl)methyl]piperazine) and molecule **8**, (1-ethyl-4-[(2-phenylthiazol-4-yl)methyl]piperazine) as well as for six oxazolidinone antibiotics that target the PTC ^21^ (Table S5) were remarkably above the 85^th^ percentile. In contrast, Molecules **1**, **3**, and **4** present low inhibition value for translation that could be explained by weak geometrical features found by the model (*see* SI Experimental).

Overall, this study shows that T_2_ relaxation is an extremely efficient and sensitive screening technique for small molecules that bind RNA. The advantage of the NMR-fragment screening approach for finding inhibitors that target the PTC is that it relies on a binding assay rather than a functional assay. It is therefore not necessary to retain the peptidyl transfer activity in the PTC fragment used for the binding assay: The effect of the binding on the peptidyl transfer activity is measured at later stages of the development when drug-sized molecules have been identified.

The *in-silico* FBVS approach enables us to reduce the number of steps in lead compound optimization.^9, 22^ Our approach for computational optimization included virtual filtration to yield drug-sized molecules containing the scaffold found by NMR, automated docking and a machine learning prediction model, which provided a fast, economic, and efficient route to two drug-sized molecules that inhibit *M. smegmatis* ribosomes.

## Conclusions

This study provides the basis for a complete design workflow that can be used for different RNA targets. By developing molecules that target the translation of mRNA by bacterial ribosomes to create proteins, we aim to inhibit the bacterial translational machinery and hence to eliminate the pathogenic bacteria. Our method, which combines experimental and computational techniques, provides proof of concept for the development of inhibitors that target the ribosomes of mycobacterial cells. Fragment-based screening complemented with virtual screening may yield false positives. However, this may be compensated by the relatively large number of hits that can be screened using more conventional *in-vitro* translation assays. Machine learning can circumvent this challenge in the future, as it can be trained directly to predict functional activity (rather than binding energy), once enough data is collected.

Machine learning provided us with an efficient prediction tool for binding energy (taking milliseconds per molecule rather than minutes on the docking simulator), allowing scanning of increasingly larger datasets of potential molecules.

The two lead compounds that we discovered could potentially be passed on to the pharmaceutical industry for testing and development into new drugs for treating active TB.

## Notes

#### Summary of Updates

Thie version of the manuscript was shortened, figures were modified and the text improved.

