## Supplemental Information for "Discovery of small-molecule inhibitors targeting the ribosomal peptidyl transferase center (PTC) of *M. tuberculosis*"

### Experimental details

#### General Experimental Information

All chemicals were obtained from Sigma, unless otherwise mentioned. Creatine phosphate and creatine kinase were obtained from Roche Molecular Biochemicals. Small molecule inhibitors were purchased from UORSY, Ukraine. All molecules were HPLC-grade pure (at least 90% pure, $H_2O$ – MeOH; Agilent 1260 Infinity systems equipped with DAD and mass-detectors and Waters Sunfire C18 prep column). Purity was verified using LC/MS (Agilent 1100 and 1200 Series system with DAD/ELSD and LC/MSD VL/SL, SL G6140A mass spectrometer) and $^{1}H$ NMR (Bruker Advance DRX 500 MHz and Varian UNITYplus 400). RNase T1 and cAMP were from Thermo Scientific. Quick calf intestinal alkaline phosphatase and T4 polynucleotide kinase were purchase from New England Biolabs, and [$\gamma$–$^{32}P$] ATP (800 Ci mmol$^{-1}$), from Perkin Elmer. The luciferase assay kit was purchased from Promega.

#### RNA preparation

RNA (GCAUCCUGGGGCUGGAGCAGGUCCCAAGG) representing the sequence of hairpin 91 in the 23S rRNA of *M. tuberculosis* (G2713-G2741) for NMR screening was obtained from Integrated DNA technology. RNA for further *in-vitro* experiments was produced in high amounts and purity as follows. The full-length rRNA of *M. tuberculosis* 23S rRNA of *M. tuberculosis* (H37Rv ATCC 27294) was cloned in a pLitmus28 vector, and a fragment of the T7 promoter–PTC (164 nt, U2646-C2809) – BSPQI (naagatc) inserted to pUC18 vector. The restriction enzyme BSPQI was used to linearize the plasmid DNA. Preparation of the DNA template for transcription, followed by *in-vitro* 'run-off' transcription and RNA purification was performed as previously described (Figure S1).[1,2] Importantly, PTC segment in different bacterial species has different nucleotide numbering: PTC sequence from: 1) *S. aureus* (164 nt, NCBI reference sequence: NR_076515.1) U2476-C2639 [UAACAGGCUUAUCUCCCCCAAG-AGUUCACAUCGACGGGGAGGUUUGGCACCUCGAUGUCGGCUCAUCGCAUCCUGGGGCUGUA-GUCGGUCCCAAGGGUUGGGCUGUUCGCCCAUUAAAGCGGUACGCGAGCUGGGUUCAGAACG-UCGUGAGACAGUUCGGUCCC]. 2) *M. tuberculosis* H37RV (NG_041982.1) U2687-C2850 [UA-ACAGGCUGAUCUUCCCCAAGAGUCCAUAUCGACGGGAUGGUUUGGCACCUCGAUGUCGGCUC-GUCGCAUCCUGGGGCUGGAGCAGGUCCCAAGGGUUGGGCUGUUCGCCCAUUAAAGCGGCAC-

GCGAGCUGGGUUUAGAACGUCGUGAGACAGUUCGGUCUC]. 3) *M. smegmatis* (NR_076104.1) U2673-C2836 [UAACAGGCUGAUCUUCCCCAAGAGUCCAUAUCGACGGGAUGGUUUGGCACCU-CGAUGUCGGCUCGUCGCAUCCUGGGGCUGGAGCAGGUCCCAAGGGUUGGGCUGUUCGCCCA-UUAAAGCGGCACGCGAGCUGGGUUUAGAACGUCGUGAGACAGUUCGGUCUC]. 4) *E. coli* (NR_076322.1) U2450-U2614 [UAACAGGCUGAUACCGCCCAAGAGUUCAUAUCGACGGCGGU-GUUUGGCACCUCGAUGUCGGCUCAUCACAUCCUGGGGCUGAAGUAGGUCCCAAGGGUAUGG-CUGUUCGCCAUUUAAAGUGGUACGCGAGCUGGGUUUAGAACGUCGUGAGACAGUUCGGUCC-CU].

RNA *in vitro* transcription was carried out in a reaction mixture containing 8 µg of linearized plasmid, 18.5 µM T7 RNA polymerase, 1 mM for ATP, CTP, GTP and UTP, 10 units of RNase inhibitor, in an appropriate reaction buffer [80 mM HEPES-KOH pH 7.5, 10 mM spermidine, 40 mM dithiothreitol (DTT), 25 mM $MgCl_2$]. Reaction mixtures (100 µL) were incubated at 37ºC for 4 h.

**$T_2$-relaxation experiment**

The nature of the binding is assessed by determining the magnetization decay rates, $R_2$, of $T_2$ relaxation-edited Carr–Purcell–Meiboom–Gill (CPMG) spectra of the fragment molecules in free form and bound to the RNA target, according to the procedure outlined by Hajduk et al.[3] Circular dichroism was performed to confirm that the RNA retain the same fold between *Mtb* hairpin 91 and the crystal structure used for the docking.

The large difference in relaxation times between the bound and unbound forms was used to filter out resonances originating from the biomolecule and the complex by using a long echo time in the CPMG experiment. In addition, the differential line broadening (DLB) for the line width at half height of the NMR peak – a measure that is indirectly related to the relaxation times – also contributes to the identification of bound fragments (Table S1).

In all samples prepared for the screening (including the initial mini-screen), the RNA concentration was 15 µM, and the concentration of the fragment molecules was 300 µM. To reduce the effect of the amide proton in amides and carbines on the $R_2$ when measuring binding of a small molecule to RNA, deuterium labeling of the RNA was used. Similarly, solutions were prepared in deuterated phosphate buffered saline (PBS), pH 7.4. All measurements were carried out at 25 °C. The molecules identified in the full screening were ranked according to the binding strength to the RNA target on the basis the NMR observables, i.e., DLB and CPMG. Some molecules gave changes in both DLB and CPMG, and others showed changes in only one observable (Table S1).

In the initial screening 10 fragment molecules from the Maybridge Ro3 1000 Diversity Fragment Library (Maybridge, Inc.) were pooled, mixed with the above-described 29mer RNA molecule that represents hairpin 91 of the *M. tuberculosis* PTC (G2754-G2782: GCAUCCUGGGGCUGGAGCAGGUCCCAAGG), 10 μM, and subjected to NMR ($T_2$ relaxation) screening. The RNA used for screening was dissolved in proton-free phosphate buffer (PBS).

#### *In silico* molecular docking

Molecular docking simulations were performed using AutoDock 4.2[4] to estimate the binding free energy ($\Delta G_{bind}$) and the poses of the investigated compounds in relation to the receptor. The PTC receptor for simulations was derived from the Cartesian coordinates of the large ribosomal SA50S originating from *S. aureus* (PDBID 4WCE) [5]. Cognate ligands chosen for this purpose were imported from the ZINC database [6]; all demonstrated at least 70% similarity with the putative inhibitor molecules suggested by the NMR experiments. The virtual screening protocol was conducted through the Raccoon implementation (http://autodock.scripps.edu/resources/raccoon), in which the receptor and ligand molecules were preprocessed for docking. The docking grid was set with 126 points in each dimension and the default spacing was 0.375 Å. The obtained grid map was centered with respect to the receptor. Free energy calculations and conformational sampling of the ligands were then carried out using the Lamarckian genetic algorithm (LGA), with an initial population size of 150 individuals, 2,500,000 free energy evaluations and 27,000 LGA generations. Clustering of the results was performed by root mean-square deviation (RMSD) calculations for the obtained poses of the ligands, with a constant tolerance of 2.0 Å. Further analyses of the results were performed using default AutoDock VS tutorial scripts [7] along with several in-house written scripts.

#### A machine learning guided approach to inhibitor design

**Data preparation:** The data contained 811 molecules with a common phenylthiazole scaffold. The remaining 107 molecules from the initial number of molecules used (919) were aligned in another coordinate system, and were therefore removed. Each molecule was composed of 20 to 51 atoms, 33 on average, 11 of which belonged to the scaffold. The raw data contained the coordinates of each atom relative to the center of the molecule. The coordinates were used to compute a total of 33 features per molecule, which are explained in Table S6. The atoms of the scaffold were numbered from 1 to 11, and for every location $i$ three features were computed (33 overall). The feature DIST$i$ is a proxy for the volume of the structure connected at location $i$ – the larger the average distance, the bigger the structure. The feature VAR$i$ is a proxy for the measure of regularity or smoothness of the structure – the smaller the variance the more regular the structure (if, for example, the variance is 0 it means that the distance between every two pairs of atoms is exactly the same; in contrast, a spiky structure results in a large variance, Figure S3). In addition to the 33 independent variables, the dependent variable for each molecule was BOND, which holds the binding energy given by the docking results.

The model features include: 1) MOD$i$, for which each position $i$ in the scaffold was designated true or false, depending on whether a chemical modification of any kind had occurred at that position, 2) DIST$i$, which represents pairwise distances within the molecule (where the pairwise distance is a proxy for the molecule's volume), and 3) VAR$i$, an integer that represents the variance of DIST$i$. The variance is a proxy for the 'regularity' of the structure, i.e., the smaller the variance, the more regular the structure (as illustrated in Figure S3). In addition to the 33 independent variables, the dependent variable for each molecule was BOND, which encompasses the calculated $\Delta$G values given by the docking procedure described in the previous section. A data matrix of 811 by 34 (denoted by *X*) was computed, where each row corresponds to a certain molecule and each column to a certain feature. Figure S4 provides a two-dimensional visualization of 400 molecules belonging to the first and fourth quartiles in terms of binding energy ($\Delta$G). The 400 data points were projected on the leading two principal components (PCs) of the covariance matrix of *X.*

We trained two statistical regression models for BOND as a function of MOD$i$, DIST$i$, and VAR$i$ – linear regression and random forest. From each model the most influential features for predicting BOND were extracted. A graphical priority map (Figure S4a) that overlays the influence of each feature with its physical location on the scaffold shows that the most influential locations, i.e., those with the highest chances of accepting chemical modification were 6 and 8. Weaker tendencies for chemical modification were found for locations 5 and 9 and a small but still significantly important tendency for 11, with the other locations not exhibiting a significant tendency for modification. Importantly, in each of the influential locations, the most significant information was given by DIST$i$ and VAR$i$ and not by MOD$i$. The larger the structure at locations 5, 8 and 9 on the phenylthiazole scaffold, the stronger the binding to the RNA, whereas, at location 8 the more regular the structure the stronger the binding to the RNA. With regard to pairwise interactions between locations, a significant link between the size of modification at location 8 and the regularity at location 6 was discovered: the larger the product $DIST_8*VAR_6$, the weaker the binding of the molecule to the RNA target. Another significant interaction was $VAR_8*DIST_8$. In this case, the larger the product, the stronger the binding. Therefore, improved binding can be obtained by modifying location 8 with a very large and a very irregular structure (quantification is presented in the *Methods* section). The pairwise interactions provide highly non-trivial constraints on the structure of the molecule, which would probably not have been revealed by "eyeballing" the data.

Based on the features we engineered molecules **1**, **3**, and **4** were predicted as lesser efficient binders. The modification at location 8 for the molecules had a structure whose size was at high percentile, but it was very irregular (at high percentile of irregularity). Another adversarial factor came from the following additional guideline that we found using machine learning: larger structure in location 8 together with an irregular structure at location 6 weaken binding. Indeed molecule **3** had an irregular structure at location 6 ($80^{th}$ percentile).

**Summary statistics:** At least one of locations 1, 2, 4, 5, 6, 8, and 9 was modified in 98% to 100% of the molecules. Locations 3, 7 and 10 were never modified. Location 11 was modified in 1% of molecules. The binding energy BOND ranged from -15.680 (strongest) to -4.77 (weakest). The average was -9.85 with standard deviation of 1.89.

**Machine learning algorithms used: a. Random forest.** Random forests constitute a well-known and widely used ensemble learning method for classification or regression that operates by constructing a multitude of regression trees at training time and outputting a value that is the average of the predictions of the individual trees. Random forests are widely used in a multitude of machine learning tasks and are considered to be among the top methods in terms of predictive power. The main advantage of random forests over linear regression is the fact that the regression surface is not linear.

We trained a random forest with 50 trees, with no restriction on the tree-depth or the number of features to be considered for each node split. The RMSE (root mean squared error) of the model prediction was 0.56 (less than third of the standard deviation of BOND).

We trained a second random forest model, this time using only the 11 MOD$i$ features. The resulting RMSE was 1.86, more than three times larger. Alongside the RMSE, a useful and commonly used statistic of the random forest is called *feature importance*. The feature importance is computed as follows: for every feature, all the splits where this feature was used as the splitting criterion are examined, and the average decrease in variance is computed (variance of the node before the split minus the pooled variance of the two children nodes after the split). The larger the difference, the more important is the feature. The features were sorted according to their feature importance value, and the values are presented in Table S7: location 8 was the most important one, then – by an order of magnitude – locations 6, 9 and 5. The remaining locations were at least another order of magnitude lower.

**b. Linear regression.** A regression model was trained, and a forward-backward stepwise procedure was used to select significant features. The significant features (with *p*-value smaller than 0.05) were (in order of significance) DIST*8*, MOD*8*, VAR*8*, DIST*5*, DIST*9* and VAR6. We then regressed BOND on those features and obtained a regression model with an adjusted $R^2$ value of 0.23; F(6,804) was 41.96 and *p*-value:< $2.2e^{-16}$. The regression coefficients are shown in Table S8.

The fact that the coefficient of all the DIST variables is negative implies that increasing their value will decrease (strengthen) BOND. Similarly the fact that VAR*8* is positive implies that increasing VAR*8* increases (weakens) BOND. Next, we regressed BOND against the six variables in the above table, and in addition all the pairwise interaction variables (e.g., VAR*6**DIST*5*). The regression adjusted $R^2$ increased to 0.25 and F(15,795)=19.11 giving p-value below 2.2e-16. Interestingly, VAR*6* became insignificant, and the product VAR*6**DIST*8* became significant. The second significant interaction was VAR*8**DIST*8* with a negative coefficient -0.9. VAR*8* and DIST*8* remained significant with coefficients 5.88 and -4.25, respectively. This means that to have a strong bonding energy, i.e., negative BOND, the structure attached at location 8 needs to be either very regular and large, or irregular and large so that the product -4.25*VAR*8**DIST*8* is much larger than 5.88*VAR*8*-4.25*DIST*8*. This is feasible since the product grows faster than the linear terms.

Source code for machine learning algorithms can be found in Github repository (https://github.com/csbarak?tab=repositories). This git repository contains also the data used for the analysis.

**_In-vitro_ translation**

The inhibition effect of compounds **1-10** and chloramphenicol as a reference compound on *M. smegmatis* ribosomes was tested in a bacterial coupled transcription/translation assay, in which the expression of luciferase gene was measured. The luciferase gene was inserted into the plasmid downstream from the T7 RNA polymerase promotor. The reaction mixture contained: 160 mM HEPES-KOH (pH 7.5), 6.5% polyethylene glycol 8000, 0.074 mg/ml tyrosine, 1.3 mM ATP, 0.86 mM for CTP, GTP and UTP, 208 mM potassium glutamate, 83 mM creatine phosphate, 28 mM ammonium acetate, 0.663 mM cAMP, 1.8 mM DTT, 0.036 mg/ml folinic acid, 0.174 mg/ml *E. coli* tRNA mix, 1 mM of each amino acid, 0.25 mg/ml creatine kinase, 0.044 mg/ml T7 RNA polymerase, *E. coli* cell-free extract, 0.003 g/l luciferase-encoding plasmid and compounds **1-10** in concentrations ranging from 152 nM to 1 mM. The effect of these compounds, at a final concentration of 300 nM, was also tested against *M. smegmatis* ribosomes. The reaction mixture was incubated at 37 °C for 1 h and terminated by the addition of erythromycin at a final concentration of 8 µM. To quantify the reaction products, luciferin assay reagent (LAR, Promega) was added at 5:3 (luciferase: reaction mix) volume ratio, and luminescence was measured. The mycobacterial reconstituted translation system yielded 40% of the signal usually obtained for the *E. coli* translation system.[8] The results were plotted (compound concentration vs. luminescence intensity) and $IC_{50}$ values were calculated.

**In line probing**

in the in-line probing assay was performed as described previously.[9] An RNA construct was prepared using T7 RNA polymerase (*see* RNA preparation). The RNA 5' phosphate was removed by quick dephosphorylation kit (New England biolabs) and phosphorylated by T4 polynucleotide kinase (New England biolabs) and [$\gamma$ -$^{32}$P]ATP (3000 Ci/mmol) (Perkin Elmer). The samples were loaded onto 10% polyacrylamide denaturing gel (8 M urea). The gel bands were detected by autoradiography, and slices containing monodispersed RNA were excised, recovered, and dissolved in ultrapure water. Labeled RNAs were incubated at 25 °C either with or without the compounds (1 mM, 250 µM, 62 µM, 16 µM) for 50 h in 10 µl reactions containing 20 mM $MgCl_2$, 100 mM KCl, and 50 mM Tris-HCl. The incubation times with RNase T1 and alkaline buffer were 2 and 5 min, respectively. The samples were dried, dissolved in 5 M urea, and loaded on 10% polyacrylamide gel. The gels were visualized using autoradiography.

**Figures S1-S4**

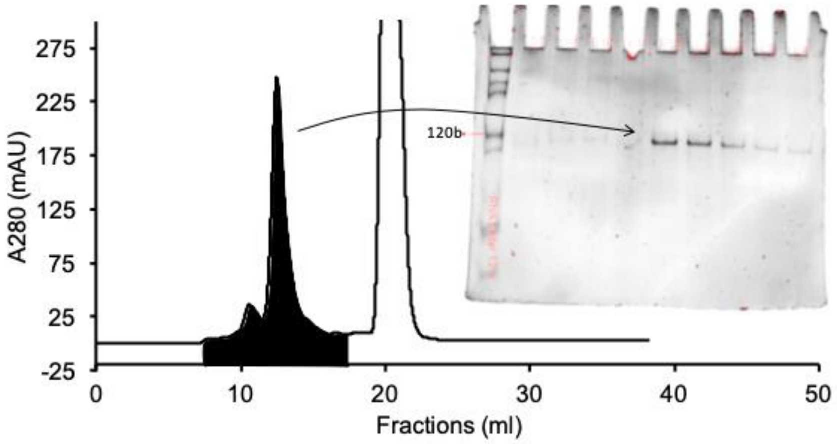

**Figure S1.** Analytical RNA sample purification. The samples were loaded into 10% TBE UREA 7M gel after size exclusion chromatography using Superdex-200 PR 26/60 column.

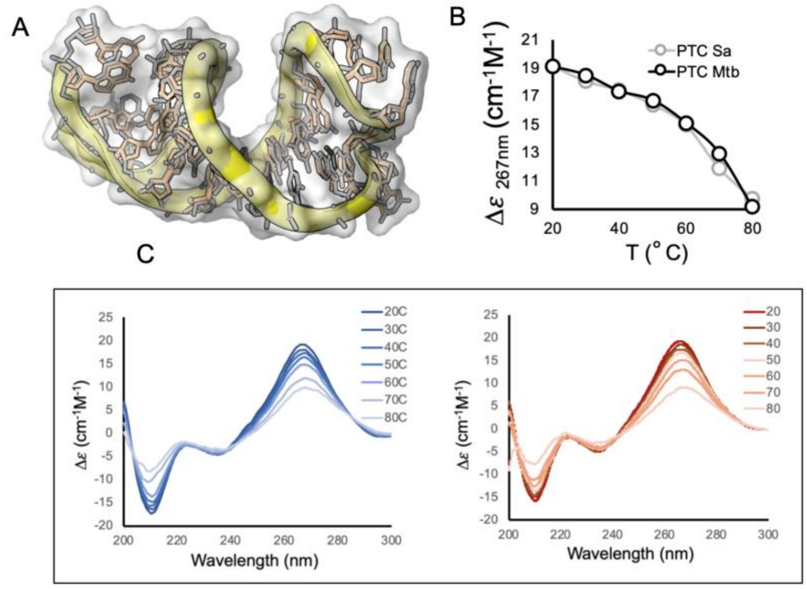

**Figure S2. Far UV CD spectra of RNA substrates** A) Short RNA hairpins that represent the fragment G2754-G2782 of the 23S rRNA of *M. tuberculosis.* B) The RNA segments (25 µM) in 20 mM Tris (pH 7.5) and 1mM $MgCl_2$ were analyzed by CD (267nm) upon change in temperature. CD spectra were measured with a 10-mm path-length cell at 25 °C. The CD spectrum is typical of an A-form RNA helix. C) Full CD spectra for the RNA from *M. tuberculosis* (left) and *S. aureus* (right)

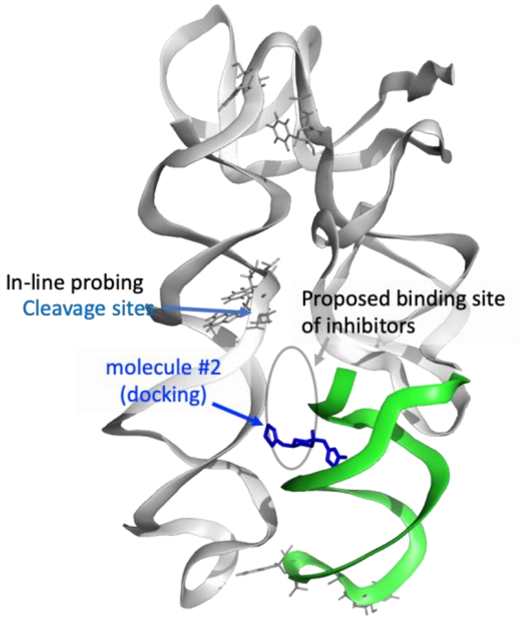

**Figure S2. The proposed binding site of inhibitors based on docking and In-line probing assay.** Molecular docking of *M. tuberculosis*-homologous *S. aureus* peptidyl transferase center (PTC) (PDB entry 4WCE) with a top-ranked resultant inhibitor (molecule **2**, marked in blue). Cleavage sites on the PTC (marked in light blue) provide an additional indication for the inhibitors' binding site (grey circle).

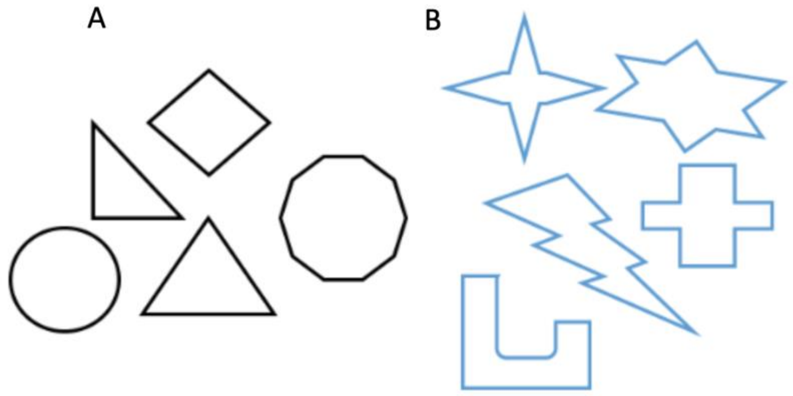

**Figure S3.** Regularity of molecular structure. The variance in pairwise distance is much smaller and therefore more regular in (A) than in (B)

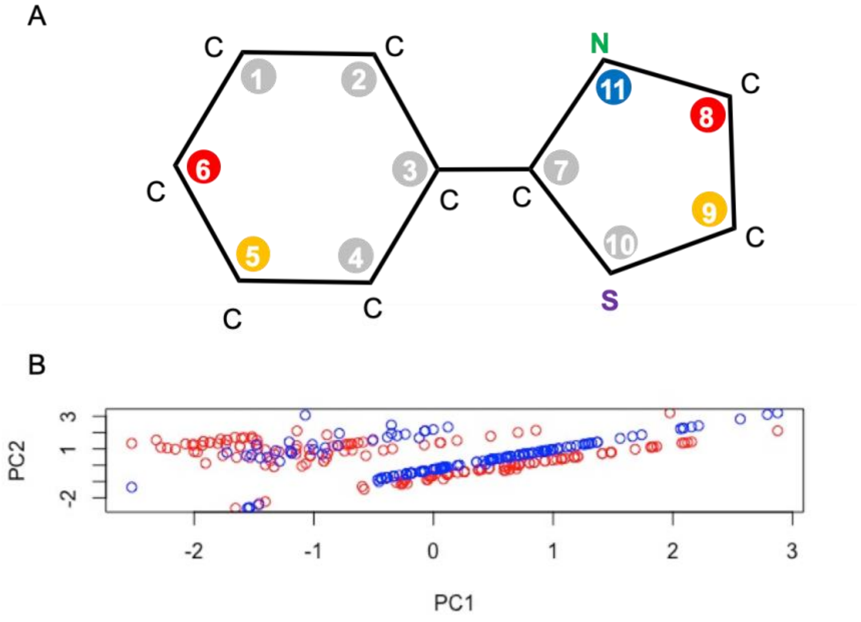

**Figure S4.** Machine learning guided structure-activity relationships. A) Each location is numbered, and its influence on the value of the bonding energy is marked with a color: red – very significant, yellow – medium significance, teal – low significance and grey not significant. B) Projection of 400 33-dimensional data points on the top two principal components of the data matrix. The 400 data points correspond to the first and fourth quartiles in terms of bonding energy. Blue points have the strongest bonding energy, and red the weakest. Clear separation can be observed between most of the blue and red points.

**Tables S1-S6**

**Table S1.** a) Summary of NMR screening results. Two binding parameters were used to determine binding affinity by using NMR fragment screening of hairpin 91 in the PTC of *M. tuberculosis*. Four fragment molecules were shown to have the best binding parameters using both Carr-Purcell-Meiboom-Gill sequence (CPMG) spectra and differential line broadening (DLB). b) molecules sorted by their binding strength to hairpin 91 of the PTC.

A

| | **Strong binding** | **Medium binding** | **Low binding** | **Total** |
|---|---|---|---|---|
| **DLB** | 10 | 12 | 9 | 31 |
| **CPMG** | 4 | 13 | 15 | 32 |
| **DLB+CPMG** | 4 | 9 | 15 | 28 |
| **Total** | 18 | 34 | 39 | 91 |

B

| | Strong binding | Mediocre binding | Low binding |
|---|---|---|---|
| differential line broadening (DLB) | (2-methylimidazo[1,2-a]pyridin-3-yl)methanol | (1-benzyl-1H-imidazol-2-yl)methanol | N-methyl-N-[3-(morpholin-4-ylmethyl)benzyl]amine |
| | (2-pyrrolidin-1-ylpyrid-4-yl)methanol | [1-(thien-2-ylmethyl)piperid-4-yl]methylamine | 2-ethyl-4-methyl-1H-imidazole |
| | [6-(diethylamino)-3-pyridinyl]methanol | 1-methyl-1H-indol-5-amine | 1-methyl-5-(trifluoromethyl)-1H-pyrazol-3-ol |
| | [3-(1H-imidazol-1-ylmethyl)phenyl]methanol | [4-(1H-imidazol-1-ylmethyl)phenyl]methanol | {2-[2-(dimethylamino)ethoxy]phenyl}methanol |
| | 2-pyrazinylmethanol | 1-(2,6-dihydroxyphenyl)ethan-1-one | N-methyl-N-[3-(2-methyl-1,3-thiazol-4-yl)benzyl]amine |
| | N-methyl-N-(2-phenoxybenzyl)amine | 4-(morpholinomethyl)aniline | 2-(2-methyl-1,3-thiazol-4-yl)acetic acid |
| | 3-(2-methyl-1H-imidazol-1-yl)propanenitrile | 3-(1H-imidazol-1-ylmethyl)aniline | methyl 3-hydroxythiophene-2-carboxylate |
| | (2-butyl-1H-imidazol-4-yl)methanol | (1-methyl-1H-imidazol-2-yl)methanol | 6-hydroxy-2,3-dihydrobenzo[b]furan-3-one |
| | [6-(1-pyrrolidinyl)-3-pyridinyl]methanol | [2-(1-pyrrolidinyl)-3-pyridinyl]methanol | [4-(morpholinomethyl)phenyl]methanol |
| | 5-methylpyridin-2-amine | (6-amino-3-pyridinyl)methanol | |
| | | 7-hydroxy-4-(trifluoromethyl)-2H-chromen-2-one | |
| | | 2-(4-benzylpiperazino)ethan-1-amine | |

| | | | |
|---|---|---|---|
| Carr-Purcell-Meiboom-Gill (CPMG) | 4-(1H-pyrrol-1-yl)benzylamine | 9-oxo-9H-fluorene-4-carboxamide | ethyl 2-amino-5-methyl-4-phenylthiophene-3-carboxylate |
| | 1-(mesitylmethyl)-1,4-diazepane | 4-(2-methyl-1,3-thiazol-4-yl)phenylamine | 3-(4-fluorophenyl)-5-(methylthio)-1H-pyrazole |
| | 4-(furan-2-yl)benzonitrile | (4-thien-2-ylphenyl)methanol | 4-benzylpiperidine-1-carboximidamide hydroiodide |
| | 4-methylpiperazinoamine dihydrochloride monohydrate | 4-(2-methyl-1,3-thiazol-4-yl)benzoic acid | 3-(2-furyl)benzonitrile |
| | | [2-(1H-pyrrol-1-yl)phenyl]methylamine | 6-ethynylquinoxaline |
| | | 4-methyl-2-quinolinol | (2-methyl-5-phenyl-3-furyl)methanol |
| | | 2-thien-2-ylaniline | N-methyl-N-(3-pyridin-3-ylbenzyl)amine |
| | | (4-phenoxyphenyl)methanol | [2-(2-furyl)phenyl]methylamine |
| | | N-methyl-N-[(1-methyl-1H-indol-5-yl)methyl]amine | N-methyl-N-(quinolin-6-ylmethyl)amine |
| | | 3-(1H-pyrrol-1-yl)benzylamine | 6-chloro-1,3-benzothiazol-2-amine |
| | | 2-(1H-imidazol-1-yl)aniline | 5-fluoro-3-(4-piperidinyl)-1,2-benzisoxazole hydrochloride |
| | | 5-chloro-2-(methylthio)aniline | 4-(2-methyl-1,3-thiazol-4-yl)benzonitrile |
| | | 4-Piperazin-1-yl-benzonitrile | 5-(4-chlorophenyl)-3-methyl-1,2,4-thiadiazole |
| | | | 5-(4-methylphenyl)-1,3-oxazole |
| | | | methyl quinoline-6-carboxylate |
| CPMG and DLB | 9H-beta-carboline | N-methyl-N-(4-thien-2-ylbenzyl)amine | N-methyl-3-(1,3-thiazol-2-yl)benzylamine |
| | 8-methyl-8-azabicyclo[3.2.1]octan-3-one oxime hydrochloride | N-methyl-4-(1,3-thiazol-2-yl)benzylamine | N-(1-benzothien-2-ylmethyl)-N-methylamine hydrochloride |
| | 7-chloro-4-piperazinoquinoline | 1,3-benzothiazol-2-ylmethanol | tetrahydrothiopyran-4-ylamine |
| | [2-(3-chlorophenyl)-1,3-thiazol-4-yl]methanamine hydrochloride monohydrate | (2-phenyl-1,3-thiazol-4-yl)methylamine | (5-phenylisoxazol-3-yl)methylamine |
| | | N-methyl-N-(3-pyridin-4-ylbenzyl)amine | N-methyl-N-(3-thien-2-ylbenzyl)amine |
| | | N-methyl-N-(3-thien-3-ylbenzyl)amine | (2-piperidinopyrid-4-yl)methanol |
| | | 6-chloro-2-(1,4-diazepan-1-yl)-1,3-benzothiazole | (1-methyl-1H-benzimidazol-2-yl)methylamine |
| | | N-methyl-N-(4-thien-3-ylbenzyl)amine | methyl 5-amino-1-benzothiophene-2-carboxylate |
| | | {2-[4-(trifluoromethyl)phenyl]-1,3-thiazol-4-yl}methylamine | 1,3-diphenyl-1H-pyrazol-5-amine |
| | | | 3-amino-6-methyl-4-(trifluoromethyl)thieno[2,3-b]pyridine-2-carbonitrile |
| | | | 4-hydroxy-2,6-dimethylbenzonitrile |
| | | | 2-methyl-4-piperazinoquinoline |
| | | | 3-piperidin-1-ylmethyl benzylamine |
| | | | 2-(3-chlorophenyl)-1,3-thiazole-4-carboxylic acid |
| | | | {3-[(4-methylpiperidino)methyl]phenyl}methanamine |

**Table S2**. Magnetization decay rates ($R_2$) determined from $T_2$–edited CPMG spectra for the four aromatic resonances of [2-(3-chlorophenyl)-1,3-thiazol-4-yl]methanamine, as indicated in Figure 2, in the absence and presence of the RNA target.

| **peak** | **T2 (us, Free)** | **T2 (us, RNA)** |
|---|---|---|
| **1** | 2500 | 167 |
| **2** | 1818 | 182 |
| **3** | 1052 | 182 |
| **4** | 2222 | 1538 |
| **5** | 1333 | 1333 |
| **6** | 1176 | 1176 |

**Table S3.** Molecular docking results for the twenty top-ranked candidate inhibitor molecules (out of 919 compounds) with *M. tuberculosis*-homologous *S. aureus* PTC. The Table shows the energetic and structural characteristics for each molecule. The ten best molecules were tested for their ability to inhibit ribosomes of *M. smegmatis.* The full list, designated as small_molecule_library.csv is available online in: https://github.com/csbarak/PTCinhibitors)

| Rank | ZINC name | Compound Number | $\Delta G_{bind}$ (kcal/mol) | Number of Atoms | Number of Torsions |
|---|---|---|---|---|---|
| **1** | ZINC37712815 | 1 | -15.68 | 21 | 6 |
| **2** | ZINC37713642_01 | 9 | -15.37 | 23 | 7 |
| **3** | ZINC37712815_01 | — | -15.33 | 22 | 6 |
| **4** | ZINC54418966 | 7 | -15.22 | 21 | 6 |
| **5** | ZINC37715345 | 5 | -15.2 | 24 | 8 |
| **6** | ZINC19595411 | 8 | -15.19 | 21 | 4 |
| **7** | ZINC23253647 | — | -15.15 | 21 | 3 |
| **8** | ZINC87590125 | — | -14.93 | 20 | 4 |
| **9** | ZINC71794395 | 6 | -14.88 | 24 | 7 |
| **10** | ZINC54418918 | — | -14.84 | 21 | 6 |
| **11** | ZINC22812775 | 2 | -14.78 | 26 | 5 |
| **12** | ZINC87590112 | — | -14.73 | 21 | 4 |
| **13** | ZINC87589974 | — | -14.66 | 21 | 4 |
| **14** | ZINC19944344 | 3 | -14.6 | 21 | 3 |
| **15** | ZINC22812775_02 | — | -14.5 | 25 | 5 |
| **16** | ZINC54418570 | — | -14.5 | 20 | 6 |
| **17** | ZINC93891293_01 | — | -14.46 | 24 | 8 |
| **18** | ZINC35154793 | 10 | -14.4 | 23 | 7 |
| **19** | ZINC35154803 | — | -14.37 | 21 | 6 |
| **20** | ZINC90290472 | 4 | -14.37 | 20 | 4 |

**Table S4.** The half maximal effective dose ($IC_{50}$) of selected candidate inhibitors.

| Compound number | IC50 (µM) |
|---|---|

| | |
|---|---|
| **1** | NA (too high) |
| **2** | 9.1 ± 1.7 |
| **3** | NA (too high) |
| **4** | NA (too high) |
| **5** | NA (too high) |
| **6** | 330.8 ± 174.9 |
| **7** | NA (too high) |
| **8** | 2.8 ± 1.4 |
| **9** | NA (too high) |
| **10** | NA (too high) |
| **Chloramphenicol** | 7.1 ± 1.9 |

**Table S5.** Predicted binding of oxazolidinone antibiotics that target the ribosomal PTC.

| | oxazolidinone | Zinc ID | Predicted ΔG binding |
|---|---|---|---|
| 1 | Eperezolid | ZINC03813328 | -11.13 |
| 2 | Linezolid | ZINC02008866 | -10.67 |
| 3 | Radezolid | ZINC40379938 | -10.86 |
| 4 | Sutezolid | ZINC03810825 | -11.12 |
| 5 | Tedizolid | ZINC43100953 | -11.28 |
| 6 | Posizolid | ZINC03982517 | -10.94 |

**Table S6.** Machine learning feature selection

| Feature name | Type | Description |
|---|---|---|
| **MOD$i$** | Binary | MOD$i$ $=1$ if location $i$ was modified, and 0 otherwise. |
| **DIST$i$** | Real | The average distance between pairs of atoms in the structure attached at location $i$. If there was no modification or it contained only one atom, the value is 0. |
| **VAR$i$** | Real | The variance of the variable DIST$i$ |

**Table S7.** Machine learning feature importance

| Feature | DIST8 | VAR8 | DIST6 | VAR6 | DIST9 | VAR9 | DIST5 |
|---|---|---|---|---|---|---|---|
| **Value** | 0.5 | 0.34 | 0.05 | 0.04 | 0.02 | 0.02 | 0.01 |

**Table S8.** Regression coefficients

| Feature | Estimate | Standard Error | p-value |
|---|---|---|---|
| **Intercept** | -9.3988 | 0.5681 | < 2e$^{-16}$ |
| **VAR6** | -1.9071 | 0.2343 | 1.50e$^{-15}$ |
| **DIST5** | -0.72 | 0.12 | 1.39e$^{-08}$ |
| **VAR8** | 1.5163 | 0.2208 | 1.30e$^{-11}$ |
| **DIST8** | -3.0386 | 0.281 | < 2e$^{-16}$ |
| **DIST9** | -1.257 | 0.2649 | 2.44e$^{-06}$ |
| **MOD8** | 6.0238 | 0.761 | 8.26e$^{-15}$ |
